# Automatic detection of spatio-temporal signalling patterns in cell collectives

**DOI:** 10.1101/2022.07.12.499734

**Authors:** Paolo Armando Gagliardi, Benjamin Grädel, Marc-Antoine Jacques, Lucien Hinderling, Pascal Ender, Andrew R. Cohen, Gerald Kastberger, Olivier Pertz, Maciej Dobrzyński

**Affiliations:** Institute of Cell Biology, University of Bern, Baltzerstrasse 4, 3012 Bern, Switzerland; Graduate School for Cellular and Biomedical Sciences, University of Bern, Switzerland; Department of Electrical and Computer Engineering, Drexel University, 3120-40 Market Street, Suite 313, Philadelphia, PA 19104, USA; Institute for Biology, Karl-Franzens-University Graz, 8010 Graz, Austria

**Keywords:** cell signalling, spatial clustering, time series, computational methods

## Abstract

An increasing experimental evidence points to physiological importance of space-time correlations in signalling of cell collectives. From wound healing to epithelial homeostasis to morphogenesis, coordinated activation of bio-molecules between cells allows the collectives to perform more complex tasks and better tackle environmental challenges. To understand this information exchange and to advance new theories of emergent phenomena, we created ARCOS, a computational method to detect and quantify collective signalling. We demonstrate ARCOS on cell and organism collectives with space-time correlations on different scales in 2D and 3D. We make a new observation that oncogenic mutations in the MAPK/ERK and PIK3CA/Akt pathways of MCF10A epithelial cells induce ERK activity waves with different size, duration, and frequency. The open-source implementations of ARCOS are available as R and Python packages, and as a plugin for napari image viewer to interactively quantify collective phenomena without prior programming experience.

## Introduction

Time-lapse fluorescence microscopy has become a routine experimental modality that has greatly furthered our understanding of cell-cell heterogeneity and the effect of signalling on cell fate determination (1–3). Live fluorescent biosensors have also revealed that signalling propagates across cell collectives; a phenomenon that is becoming another important organisational principle in biology. Even though we are only beginning to understand the rules that govern the emergence of spatial correlations from single-cell signalling, their functional significance has been reported in various systems. Notably, the coordinated activation of extracellular signalregulated kinase (ERK) in spatial clusters of cells plays an important role in the maintenance of epithelial homeostasis (4–6), acinar morphogenesis (7), osteoblast regeneration (8), cell cycle progression (9), and in the coordination of collective cell migration (10, 11). Calcium waves observed in 2D monolayers trigger and facilitate the extrusion of oncogenically transformed cells from the epithelium, thus also contributing to its homeostasis (12).

Waves rely on an active process in which individuals relay the signal from their neighbours (13). They are an important class of self-organising behaviours that have emerged repeatedly at different biological scales, arguably because they can robustly transmit information at long distances throughout a community (14). At the cellular level, voltage-gated sodium channels mediate the action potential that propagates in neurons (15). A rapid initial development phase in large animal eggs requires long-distance coordination of mitotic events, which can only be achieved by signalling waves (16). At the cell population level, waves rely on cell-cell communication as is the case of cyclic adenosine monophosphate (cAMP) waves in *D. discoideum* (17), calcium waves in animal development, adult animal tissues and plants (18–20), depolarisation waves in the heart that control heartbeat (21), mechanical waves (22), and genetic waves in the presomitic mesoderm (23). Wave patterns also occur in communities of independent multicellular organisms. Giant honeybees flip their abdomens in a coordinated fashion that results in shimmering waves that repel an intruder that endangers the colony (24).

Despite waves being so common, a general and an easy-touse computational tool to identify them and extract useful statistics for further analysis is still missing. Notably, the fields of geographic information science and ecology have accumulated a multitude of statistical methods to analyse space-time patterns in weather prediction, animal migration, and voting preferences. Here, we build upon these methods and develop a computational tool for the Automatic Recognition of COllective Signalling (ARCOS) events and their characterisation with various statistics. The open-source code is freely available as R and Python packages, along with extensive documentation. We also developed a plugin for Napari – a fast, interactive, multidimensional image viewer for Python (25). The plugin provides a graphical interface to ARCOS and facilitates an intuitive discovery environment to quantify collective phenomena in time-lapse images.

In the following sections, we explain the ARCOS algorithm and apply it to several biological systems. First, we used an optogenetic actuator that activates the MAPK network to generate synthetic waves of ERK activity *in vitro* in an MCF10A epithelial monolayer. This way we could establish a ground truth to test the algorithm in a controlled setting. We then studied mammary epithelial cells expressing different oncogenic mutations. We found that ERK activity waves exhibit mutation-specific features, suggesting different alterations of intercellular communication. Further, we quantified apoptosis-induced waves of ERK activity, calcium waves in renal cells, and collective shimmering in a colony of giant honeybees. Finally, we tackled 3D geometry and identified ERK waves in mammary acini *in vitro*.

## Results

### The algorithm

To explicate the core ARCOS algorithm, let us consider a radially expanding wave, a “firework”, as shown in Fig. 1A, where we schematically depict 3 consecutive frames of a developing activation cluster. Such events have been observed, for example, in epithelial monolayers where a correlated activation of ERK is typically initiated by an apoptotic cell (4–6). In the 1^st^ step, we apply a spatial clustering algorithm *dbscan* (26) to every frame of the time-lapse to identify pockets of active cells. The clustering requires two parameters: the search radius *ε* and the minimum number of points *minPts* within that radius. We recommend setting *ε* to ≈ 2 distances to the first nearest neighbours, and *minPts* to the dimensionality of the problem plus one, i.e., 3 for 2D, 4 for 3D, etc. (26).

**Fig. 1.**
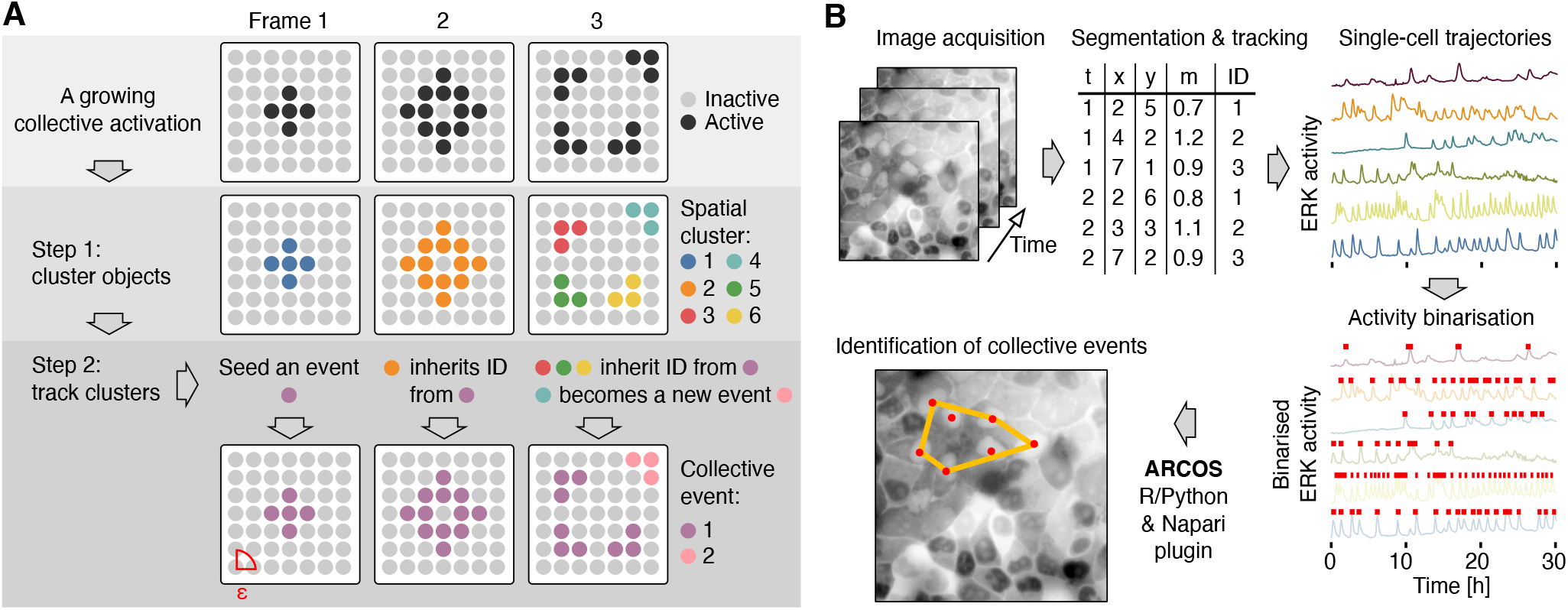
(A) Demonstration of the ARCOS algorithm on a growing activity cluster. Protein activity of cells arranged in a mesh can assume two states, inactive (light grey) and active (black). In step 1, a *dbscan* algorithm spatially clusters active cells independently in each frame. In step 2, clusters are linked between frames. The cluster in frame 1 forms a seed of a collective event. The cluster in frame 2 is linked to this seed cluster because several of its member cells are within the neighbourhood radius *ε*. In frame 3, only clusters #3, #5, & #6 are linked to the previous frame’s cluster. Cells in cluster #4 are too far and thus form a new seed of a collective event. (B) Image acquisition and analysis pipeline. Segmentation & tracking software identifies cells in multichannel time-lapse movies and produces a long-format output where every row is the measurement *m* in a cell *ID* at coordinates (*x, y*), at time *t*. Single-cell protein activity time series are then detrended and binarised. Binary activity time series are then fed into ARCOS to identify collective activity.

In the 2^nd^ step, clusters identified in the 1^st^ step are linked between frames based on their relative distance. The 1^st^ time frame contains only a single 5-cell spatial cluster #1 that becomes the seed of a new collective event. The 2^nd^ frame also contains a single but a bigger cluster #2. To link clusters #1 and #2, we calculate the nearest neighbour distance between cells that comprise both clusters. In our case, at least one cell from cluster #2 is within the *ε* distance to cells in cluster #1. Hence, cells in cluster #2 inherit the cluster identifier of the seed cluster #1. The 3^rd^ frame contains 4 spatial clusters #3-#6. Based on nearest neighbour distances between these cells and cells from the cluster in the previous frame, we conclude that only clusters #3, #5 and #6 can be linked to the growing collective event from the previous frame. Cells from cluster #4 are farther than *ε*, therefore they become a seed of a new collective event. Fig. S1A outlines the algorithm and Fig. S1B demonstrate identification of multiple collective events in a synthetic time-lapse.

A sample workflow to identify collective ERK activation in a 2D MCF10A epithelium can be summarised as follows (Fig. 1B). After acquiring multichannel time-lapse movies, segment the images and track cells over time. The resulting single-cell ERK activity time series form a space-time pattern (*x, y, m, t*), where (*x, y*) are the centroid coordinates of nuclei, *m* is the measurement of ERK activity, and *t* corresponds to time. The input for ARCOS should only contain “active cells”, i.e., spatial coordinates and time, (*x, y, t*), of cells with the measurement *m* greater than a threshold. Therefore, a crucial step in the pipeline involves binarisation of the single-cell activity dynamics.

We implemented several methods to binarise time series data in ARCOS. The most straightforward approach applies a fixed threshold to the measurement. However, data from time-lapse microscopy often exhibit high variability in the baseline due to different expression levels of the biosensor. Additionally, high intensity light and long exposures can lead to photo-bleaching of fluorescent probes, which may cause long-term trends. To address this, ARCOS offers a detrending method based on a running median or on a fit to a polynomial (Fig. S1C and Materials & Methods). An alternative approach to measurement binarisation utilises local indicators of spatial association (LISAs), which are a class of statistics that quantify local spatial correlation and clustering (27). A LISA statistic applied to individual time frames of the spacetime pattern (*x, y, m, t*) yields a coefficient of local spatial association *ρ* at every coordinate (*x, y*). For our purposes, we are looking for pockets where the coefficient is high, which would correspond to regions with high and positively correlated single-cell ERK activity. After thresholding *ρ* we obtain a space-time pattern (*x, y, t*) that is restricted only to cells that form pockets of high activity (Fig. S1D). We assume these clusters estimate the location of active cells, and we use this subset as an input for ARCOS.

The choice of the binarisation algorithm depends on the dataset and should be evaluated on a case-by-case basis. The main disadvantage of the approach based on smoothing and detrending is that it requires long time series to apply the long-term detrending filter to reliably estimate the baseline. The recommended window for such a filter usually amounts to the quarter of the time series’ length. However, this condition may be difficult to satisfy due to image segmentation errors that give rise to short, single-cell time series. Contrary to the former method, LISA-based binarisation is not contingent on long time series because objects are analysed independently in individual frames. This lax requirements would make it a preferred approach, however we noticed that LISAs are better at detecting larger, gradually developing events.

### Optogenetically induced collective ERK activation

Aside from synthetic datasets used during the algorithm development, we also wanted to test the code on a biological system, with the actual variability in protein activity, but in a controlled setting. To this end, we utilised an MCF10A cell line with a stably expressed optogenetic actuator-biosensor system described earlier (10) (Fig. 2A). Using an optoRAF actuator, we could induce ERK activity via the RAF-MEKERK pathway in individual cells with light pulses and measure ERK activity using the cytoplasmic/nuclear translocation of the ERK-KTR biosensor. Additionally, we used a digital mirror device (DMD) to stimulate desired regions of the epithelium over time, thus generating synthetic collective ERK activity patterns that would mimic various aspects of spontaneous waves (Fig. 2B). Notably, the ERK activity induced by optoRAF did not propagate to neighbouring cells, therefore we could precisely control the propagation of synthetic patterns.

**Fig. 2.**
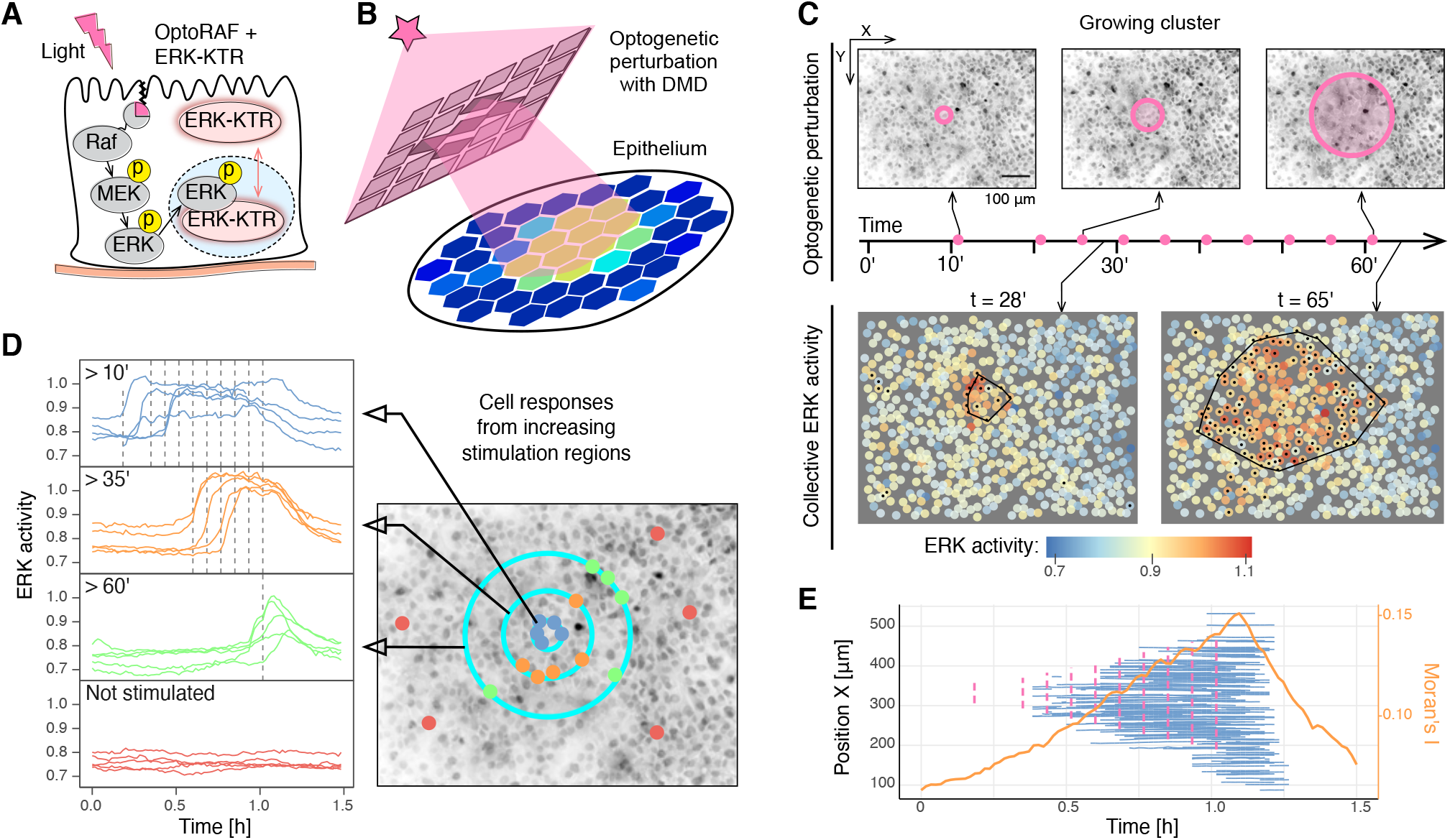
(A) Optogenetic actuator/biosensor circuit comprises the optogenetic actuator optoRAF tagged with mCitrine, the ERK biosensor (ERK-KTR) tagged with mRuby2, and a nuclear marker (H2B) tagged with miRFP703. (B) Schematic of a spatially-constrained perturbation of the 2D epithelium with a digital mirror device (DMD). (C) Repeated, pulsed optogenetic perturbation of the MCF10A WT epithelium with a growing circular region using DMD. Upper row: inverted raw images from selected time points. The circle outlines the stimulation region. Pink dots on the timeline indicate the timing of a 100 ms-long optogenetic pulse. Lower row: quantification of ERK activity from image segmentation for selected time points. Black dots indicate active cells identified from detrended and binarised ERK activity time series; black polygons indicate a convex hull around collective activation. (D) Single-cell ERK activity in different regions of the field of view. Vertical dashed lines indicate the timing of the optogenetic stimulation. Cells in the centre are stimulated at all times after 10’; cells farther away are stimulated only with pulses that occur after 35’, etc. (E) Spaghetti plot of collective ERK activation: an X-axis projection of single cells from panel C during their participation in a collective event. Vertical dashed lines indicate the timing and position of the optogenetic stimulation. Orange solid line is the Moran’s *I* measure of spatial autocorrelation calculated from single-cell ERK activity data.

In the first scenario, we repeatedly stimulated a growing circular region of the epithelium at 5’ intervals, starting at 11’, with 100 ms light pulses. We observed a clear ERK activation in the stimulated regions (Fig. 2C, top row). As expected, our algorithm applied to detrended and binarised ERK activity time series identified a single activity cluster that grew over time (Fig. 2C, bottom row and Video S1). To verify that the optogenetic stimulation was confined to desired regions, we inspected single-cell ERK activity time series (Fig. 2D). Cells from the innermost region received all stimulation pulses after the 10’ mark. ERK activity increased sharply, although several responses were delayed and ensued only after the 2^nd^ or even 3^rd^ pulse. We attribute this delay to biological variability and the fact that the cells are at the border of an already small activation region. Cells farther away from the centre responded accordingly to pulses that they were exposed to, although several cells were activated earlier, likely by the illumination of regions with smaller radii. The ERK activity in cells not exposed to DMD illumination remained at the baseline.

We concluded that our protocol was sufficiently selective to control the stimulation of a growing ERK activation cluster, and that the ARCOS algorithm correctly identified such a synthetic wave. An intuitive visualisation of that result is shown in Fig. 2E in a “spaghetti” plot. Therein, the spatiotemporal evolution of collective ERK activity was projected onto the 2D (*x, t*) plane. Each trace corresponds to a single cell involved in a collective event identified by the algorithm. We confirmed that the collective ERK activation grew over time and within the bounds of the DMD illumination. As a cross-check, we compared the result obtained from ARCOS with an independent quantification of the global spatial correlation per frame using Moran’s *I* statistic (28). The statistic is commonly used in the field of geographic information science to quantify spatial phenomena. The Moran’s *I* coefficient increased with the growing collective ERK activity region and peaked when that region was the biggest (Fig. 2E). This result confirmed a spatially correlated cluster as detected by our algorithm.

Subsequently, we tried other illumination scenarios to further challenge the algorithm. To test the tracking, we illuminated a pattern with a circle that moved around the field of view (Fig. S2A,B and Video S2). Another common scenario observed in the epithelium prompted us to apply a circular pattern that was slowly splitting into two detached regions (Fig. S2C,D and Video S3). In both cases, our algorithm correctly identified the spatio-temporal progression of the optogenetically-induced ERK activity region. Due to a longer, 2h 40’, acquisition time, we detected additional ERK activity clusters that stemmed from apoptotic events and/or a spontaneous activation. The results from the optogenetic experiments assured us that our image analysis pipeline performed well in a controlled setting with biological variability, and that we could apply it to more challenging scenarios.

### ERK waves in MCF10A mutants

One of our earlier studies demonstrated that apoptosis of single cells within an epithelium triggers waves of ERK activity pulses of constant width and amplitude (5). These ERK waves act as a survival signal that protects 2-3 layers of neighbouring cells from further apoptosis, thus contributing to epithelial homeostasis. The ERK waves are trigger waves in which the extruding apoptotic cell activates matrix metalloprotease (MMP)-mediated cleavage of pro-EGFR ligands. Subsequently, the epidermal growth factor receptor (EGFR) activates ERK in the adjacent cells (4, 10). ERK-mediated activation of myosin contractility might then provide a mechanical input that activates ERK in farther cells through MMP/EGFR signalling. This explains how the ERK wave emerges through a mechanochemical feedback loop (11). We showed that in starved conditions, the majority (67%, SD=7%) of such waves resulted from individual apoptotic events in WT MCF10A cells. Given those past results, we were curious how oncogenic mutations that affect the MAPK network might influence the spatio-temporal ERK activation. Therefore, we explored ERK waves in starved MCF10A cells with oncogenic heterozygous mutations, KRAS^G12V^ or PIK3CA^H1047R^, that activate the MAPK/ERK pathway through different mechanisms and frequently occur in cancer. In the KRAS^G12V^ mutant, loss of GTP hydrolysis leads to increased activation of the RAF kinase, thus activating MEK and ERK. This mutation also increases the secretion of amphiregulin (29), that by binding to EGFR further activates ERK. The H1047R mutation in PIK3CA, the alpha subunit of PI3K, intermittently activates the non-mutated MAPK/ERK pathway via EGFR also by amplifying amphiregulin expression/secretion (30, 31).

As expected, at the population level, in starved conditions, without external stimulation, the average ERK activity was higher in the mutants than in the WT line (Fig. 3A). Notably, the increased basal activity occurs without ERK activity waves due to apoptosis, that is negligible in mutants. At the single-cell level, ERK activity still exhibited nonperiodic pulses, although their number and shape markedly differed between the two mutant cell lines (Fig. 3B). WT cells exhibited sharp, isolated and rare ERK pulses as observed earlier (2). In the KRAS^G12V^ mutant cells, individual ERK pulses were still discernible, but often lasted longer and formed “massifs” of ERK activity. In the PIK3CA^H1047R^ mutant cells, as already reported (7), ERK pulses retained the same shape, duration, and amplitude as in WT cells, but the frequency of these ERK pulses increased.

**Fig. 3.**
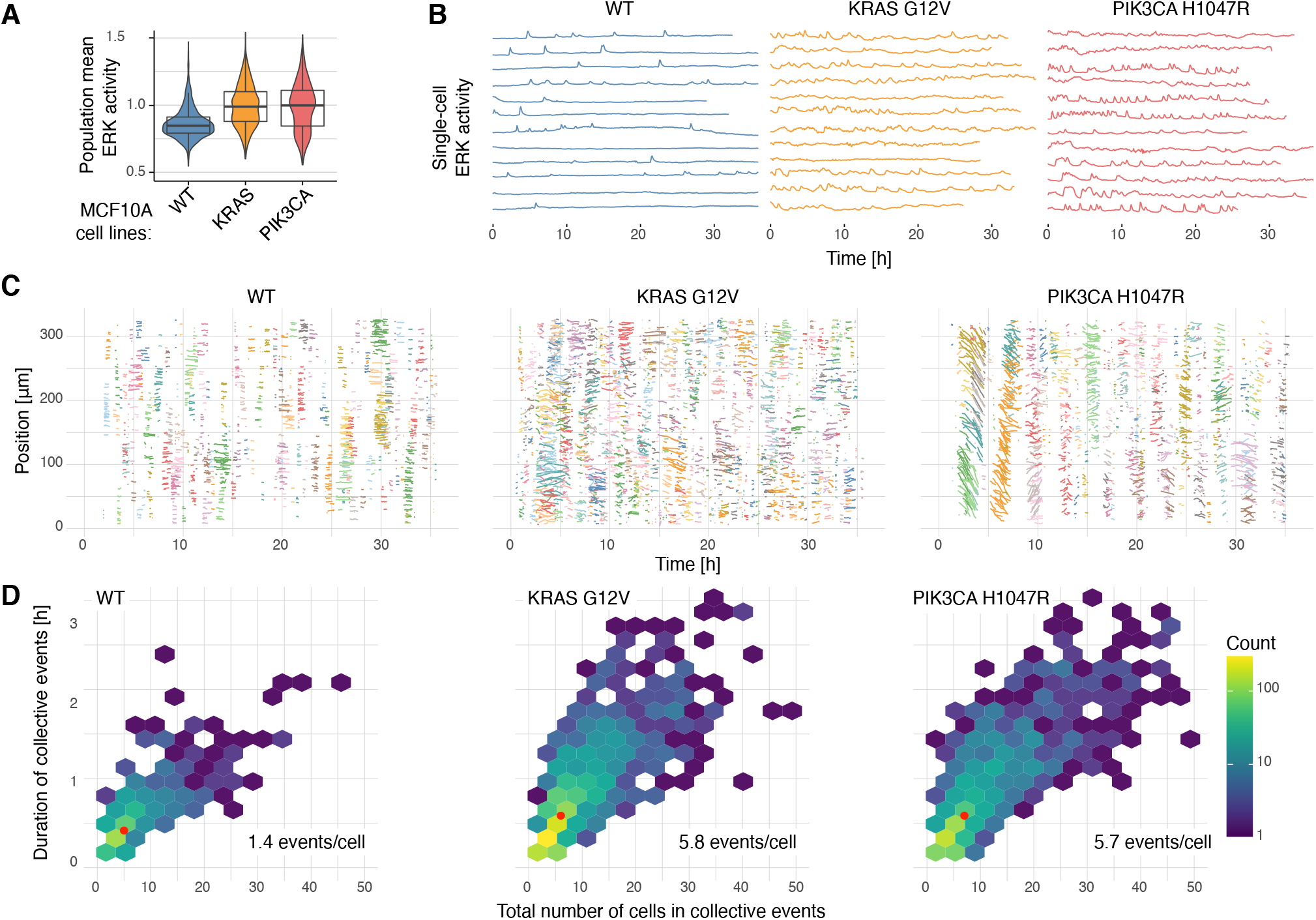
Quantification of ERK activity in starved MCF10A WT, and KRAS^G12V^ and PIK3CA^H1047R^ mutant cell lines. (A) Population-averaged ERK activity at 10h after the start of acquisition. Box plots depict the median and the 25^th^ and 75^th^ percentiles, whiskers correspond to minimum and maximum non-outlier values; *N ≈* 1300 cells per cell line. Random sample of single-cell ERK activity dynamics. (C) X-axis projections of single cells during their participation in collective events. Events are coloured using a finite discrete palette, thus colours repeat over time. Data from single chosen FOVs. (D) Hex-bin plots of the total number of cells involved in collective events vs. their duration. Each hexagon is coloured according to the number of collective events; note the logarithmic scale of the colour bar. Red dots indicate median statistics. Data pooled from 4 (WT; 489 events), 5 (KRAS^G12V^; 1583), and 6 (PIK3CA^H1047R^; 1116) FOVs.

We then wondered how different single-cell ERK activity dynamics translated into spatial correlations in the epithelium. Because the cell size and hence the epithelial density varied between the cell lines, we set the *ε* radius for cluster identification to 1.5 times the median 1^st^ nearest-neighbour distance between active cells, i.e., 26 *µ*m for WT and KRAS^G12V^, and 33 *µ*m for PIK3CA^H1047R^. The resulting spaghetti plots generated by ARCOS (Fig. 3C) provide an overview of collective events during the entire acquisition in individual fields of view (FOVs). ERK activity was the least spatially correlated in WT cells where waves were sparse, had a median diameter of 50 *µ*m (2-3 cell layers) and were usually triggered by apoptotic events. In comparison, ERK waves in KRAS^G12V^ were more correlated; the events were longer (up to 2h), bigger (median diameter 56 *µ*m) and more frequent. We observed even more spatio-temporal correlation in the PIK3CA^H1047R^ mutant. Notably, during the first 10h of acquisition we saw massive, up to 150 *µ*m in diameter, waves of ERK activity. The latter occurred every 2-3 hours, spanned the entire FOV and coincided with large-scale deformations of the epithelium as seen in Video S4 and reflected by angled segments in the spaghetti plot, indicative of rapid cell motility events.

Given the information about individual waves, we then quantified their duration and size, i.e., the total number of involved cells. We depict this relationship in the hex-bin plot in Fig. 3D which avoids overplotting by creating hexagonal binning tiles that are coloured according to the number of points in those bins. Even though the median statistics were similar across the cell lines, the distributions varied and confirmed our observations from spaghetti plots: small and short-lasting ERK activity waves in WT cells, and bigger and longer-lasting events in the mutants. The number of events normalised to the average number of cells in the FOV during the acquisition revealed that the mutants triggered 4 times more collective events compared to the WT (5.8 vs 1.4 events/cell). Finally, we investigated the evolution of collective events by looking at the displacement of their centroids. A firework-like collective ERK activation centred around an apoptotic extrusion as observed in WT cells exhibited a small displacement of around 25 *µ*m, which amounts to a single cell layer (Fig. S3). We recorded much larger centroid displacements in mutant cells, which partly stem from an increased cell migration but also reflect the travelling nature of collective waves, especially in the case of the PIK3CA^H1047R^ cells.

So far, the comparative analysis of 3 cell lines demonstrated how much new information, beyond population averages or even single-cell dynamics, our approach can extract. Even though ERK activity was equally elevated in the mutants, the mutations affected the space-time distribution of ERK activity pockets in different ways. From small and shortlived fireworks in WT cells to larger and longer-lasting waves in KRAS^G12V^ to massive pulsatile waves in PIK3CA^H1047R^, each cell line had a signature ERK wave behaviour with different size, duration, frequency, and displacement. In the next sections, we apply ARCOS to other systems to further demonstrate the versatility of the method and its ability to quantify space-time phenomena at different space-time scales.

### ERK waves in the NRK-52E epithelium

We quantified ERK waves in the NRK-52E renal epithelial cells that express the EKAREV-NLS ERK sensor. The majority of the waves in this cell line are triggered by apoptosis, as reported earlier (32). Since waves are bigger and isolated from each other, we used a LISA-based binarisation of ERK activity as the pre-processing step for ARCOS. To identify pockets of spatial dependence, we calculated the local *G*^*∗*^ statistic – a type of LISA measure that is particularly suited for that task (33). We used R implementation available in the elsa package (34) and applied it to individual frames of the (*x, y, m, t*) spacetime pattern from image segmentation. After thresholding the resulting statistic, we obtained the (*x, y, t*) pattern only with active cells, which we then used in ARCOS (Fig. 4A and Video S5).

**Fig. 4.**
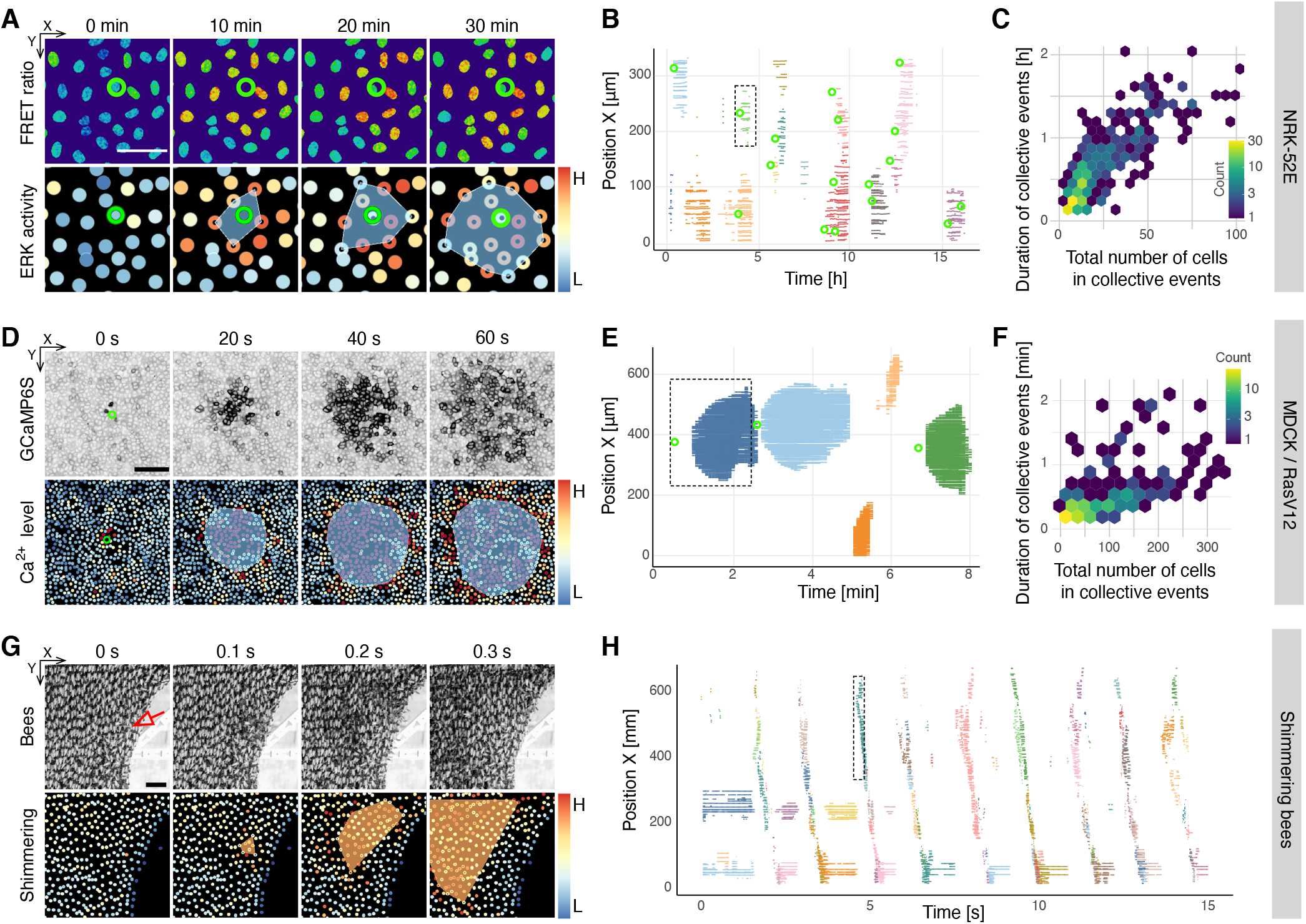
ERK activity waves in the NRK-52E epithelium (A-C), calcium waves in the MDCK epithelium (D-F), and shimmering bees (G,H). (A) Snapshots of a single ERK wave in starved NRK-52E EKAREV-NLS cells induced by the apoptotic cell indicated by a green ring. Top row shows FRET ratio calculated pixel-by-pixel from donor and acceptor images. The bottom row shows nuclei coloured by ERK activity and a convex hull of the collective event calculated by ARCOS. Black dots indicate active cells calculated from the binarisation step with the *G*^*∗*^ statistic. Scale bar, 50 *µ*m. (B) A spaghetti plot of the space-time propagation of collective events in the NRK-52E epithelium. Overlaid as green rings are manually annotated apoptotic events. (C,F) Hex-bin plot of the duration vs. the total number of cells involved in a single collective event. Hexagonal tiles are coloured according to the number of events in that bin. Note the logarithmic scale of the colour bar. (D) Snapshots of a single calcium wave and ARCOS quantification in the MDCK epithelium with the apically extruded cell indicated with a green circle. MDCK epithelial cells stably express GCaMP6S – a GFP-based intracellular calcium sensor. Scale bar, 100 *µ*m. (E) A spaghetti plot of the space-time propagation of collective events in the MDCK epithelium. Overlaid as green rings are manually annotated RasV12-expressing cells that undergo apical extrusion triggered by calcium waves. (G) Snapshots of a single shimmering bee wave induced by a dummy wasp presented to the colony on the right of the field of view. A leader bee that initiates the wave is indicated with a red arrow. The bottom row shows bees coloured by the amount of their abdomen flipping and a convex hull of the collective event identified with ARCOS. Scale bar, 50 mm. (H) A spaghetti plot of the space-time propagation of bee shimmering waves from the full time-lapse. (B,E,H) Dashed rectangles indicate space-time sections from panels A, D, G.

The spaghetti plot confirms that apoptotic events trigger ERK activity waves (Fig. 4B). To corroborate this result with an independent method, we calculated Moran’s *I* – a global indicator of spatial correlation for individual frames as in Fig. 2E. We usually observe 1-2 events at a time in the FOV. Hence, we expect the global correlation statistic calculated for the entire FOV to peak at the time of increased spatial correlation due to collective ERK activation. Indeed, in Fig. S4A we confirm that Moran’s *I* peaks at the time of collective events identified with ARCOS. After pooling 340 events from 11 fields of view, we find that the majority of events last up to 1 hour and involve up to 50 cells. Thus, the events are shorter-lived but bigger compared to waves observed in all MCF10A cell lines analysed before (Fig. 4C).

### Calcium waves in the MDCK epithelium

We then tested ARCOS on a different cell line and biosensor by analysing calcium waves that trigger apical extrusion in the MDCK epithelium (12) (imaging data, courtesy of Yasuyuki Fujita). Apical cell extrusion is an important homeostatic mechanism that eradicates harmful or suboptimal cells from the epithelium. In the experiment, the Myc-RasV12 oncogene was transiently expressed mosaically within a monolayer of MDCK cells. Due to their harmful potential, RasV12transformed cells were apically extruded from the epithelium. Notably, before the extrusion, calcium levels acutely increased in RasV12 cells and radially propagated to neighbouring cells. Such waves have been shown to trigger and facilitate the process of cell extrusion.

Again, due to the isolated nature of the waves and their apparent large size, we used the *G*^*∗*^ statistic to identify pockets of active cells (Fig. 4D,E and Video S6). After pooling 200 events from a single, 9-hour long time-lapse, we find that a single collective calcium wave propagates radially from the extruded cell, can involve up to 300 cells, and can last up to 2 minutes (Fig. 4F). Spatial correlation calculated for each frame using global Moran’s *I* statistic peaks when calcium waves occur, which further strengthens our identification of collective events with ARCOS (Fig. S4B).

### Shimmering bees

Even though we developed ARCOS to tackle collective phenomena in cell signalling, our image/data processing pipeline can also quantify spatiotemporal phenomena in the context of ecology. As a demonstration, we analysed a previously published timelapse dataset of a giant honeybee colony (*Apis dorsata*) that collectively responds to a computer-controlled dummy wasp hovering in front of the honeybee nest (35). The giant honeybees responded to the wasp by flipping their abdomens in a simultaneous and cascadic way, generating a wave-like visual signal for external observers. The visual effect achieved by this repetitive collective response aims to repel the threat (36, 37). Pulses connected to these visual patterns at the nest surface also lead to vibrations within the nest, and these are suitable for informing nest mates, even on both sides of the nest, about the threat status (24, 35, 38–41).

As described above, shimmering waves propagate across the surface of the giant honeybee nest in response to a repeated exposure to the dummy wasp presented to the colony on the right side of the FOV (Fig. 4G). Notably, our quantification confirms an interesting aspect of the shimmering phenomenon. Each vertical streak in the spaghetti plot (Fig. 4H) corresponds to a wave that propagates from right to left of the FOV. However, we note that each of such streaks actually consists of several “sub-waves” as indicated by different colours within the streak. Such a shimmering wave is typically initiated by “leader” bees on the nest surface, assembled in so-called trigger centres across the nest surface (24, 38, 42). A closer inspection of the Video S7 confirms our observation that a single exposure to the dummy wasp induces several smaller waves consecutively triggered by the leader bees scattered across the colony. These trigger centres were primarily arranged in the close periphery of the mouth zone of the nest, around those parts where the main locomotory activity occurs throughout daytime.

### ERK waves in 3D acinus

Previously, we reported waves of ERK activity during the morphogenetic program of 3D MCF10A acini *in vitro* (7). One crucial stage of this process is the formation of a lumen by apoptosis of inner cells. We found that ERK waves coordinate spatial separation into two domains: the outer cell population, that survives, and the inner cell population, that undergoes apoptosis. Since the ARCOS algorithm can handle any integer dimensionality, we were curious to analyse ERK activity waves in this system. The top row of Fig. 5A shows a single wave in a maximum projection of the raw ERK-KTR channel. This example collective event starts at the surface of the acinus and propagates over time to other cells in the outer layer (Fig. 5B and Video S8). Only towards the end, some inner cells are participating in the event. This observation reflects a more general trend that we reported earlier based on pooled data from 11 acini (7). The majority of collective events initiate and propagate at the outer layer of the acinus, thus providing a survival signal to those cells. The inner domain exhibited less spontaneous ERK pulses and also received fewer pulses due to waves that remained predominantly in the outer layer. Such a spatio-temporal distribution of waves creates a lower survival signal in the inner domain, which contributes to a higher apoptosis rate and clearing of the lumen. The formation of hollow acini recapitulates key features of *in vivo* breast alveoli and makes this system an attractive model to study the effect of MAPK signalling on lumen morphogenesis.

**Fig. 5.**
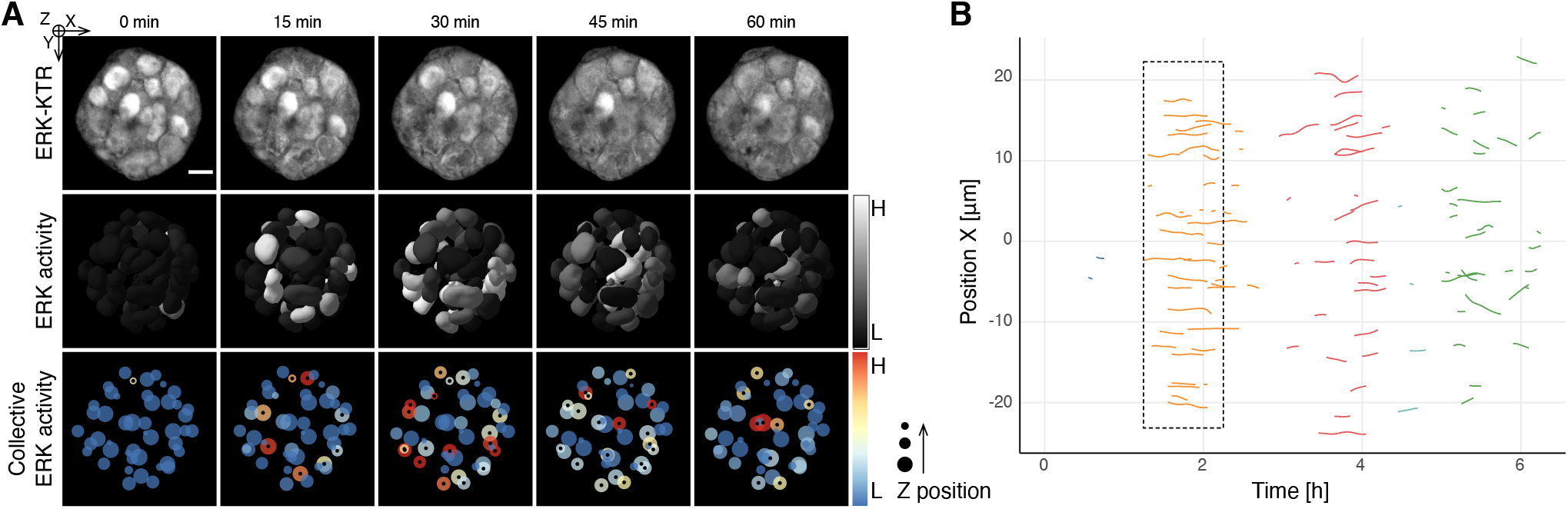
Quantification of collective ERK activation in a 7-day old MCF10A WT acinus. (A) Snapshots of a single collective event. Top row – maximum projection of the ERK-KTR channel; middle row – cell nuclei shaded according to detrended & normalised ERK activity; bottom row – nuclear centroids coloured by ERK activity and sized according to *z* distance. Black dots indicate active cells. Scale bar, 20 *µ*m. (B) Spaghetti plot of the space-time propagation of several collective events during the entire acquisition. The dashed rectangle indicates the space-time section from panel A.

## Discussion

We introduced a novel tool for detecting and quantifying collective spatio-temporal phenomena with focus on cell signalling. We validated our algorithm on several biological systems with drastically different spatio-temporal collective behaviours. We analysed ERK activity waves in an epithelial monolayer that propagate radially on the timescale of 10-30 minutes, radial calcium waves on the timescale of 10-30 seconds, directional waves in a shimmering bee colony and in a 3D culture model of mammary acini. The open-source code provided as R and Python packages can be used in Jupyter notebooks or in batch processing pipelines. The Napari plugin enables anyone without extensive programming knowledge to explore parameters through an intuitive GUI on a platform that emerges as a de-facto standard for viewing multidimensional images (Fig. S5 and a screencast with a demonstration of the plugin in Video S9).

### ARCOS as an exploration tool

We envision ARCOS as a framework to explore emergent community effects and spatio-temporal phenomena in cell communities and ecosystems. Quantification with our tool will help unravel rules that govern the dynamics of collective processes that emerge from single cells. For example, several intriguing insights become apparent from our comparison of collective signalling between WT and oncogenic cell lines. By focusing the analysis from population averages to emergent spatio-temporal phenomena of cell ecosystems, we revealed a new layer of MAPK signalling complexity. We show that both mutations augment the population-averaged ERK activity (Fig. 3A) by altering the single-cell and collective ERK dynamics. Compared to WT, both mutant cell lines exhibited 4 times more collective events per unit of time that lasted longer and involved more cells (Fig. 3D). However, ERK waves in the KRAS^G12V^ cells were more randomly distributed compared to timely, recurring events in the PIK3CA^H1047R^ mutant (Fig. 3C). We hypothesise the following mechanisms behind this observation.

KRAS^G12V^ cells have a higher ERK activity baseline and are unable to rapidly switch off ERK due to increased Raf input that overrides several negative feedbacks of the MAPK pathway (43). The resulting prolonged ERK activity periods feed into the mechanochemical feedback loop that promotes the formation of ERK waves and propagates them without apoptotic cells. An increased expression/secretion of amphiregulin might enhance this process even more (29). A different mechanism drives the dynamics of collective events in the PIK3CA^H1047R^ mutant, even though the events had a similar size/duration distribution to the KRAS^G12V^ mutant (Fig. 3D). The excitable MAPK pathway with intact negative feedbacks processes intermittent inputs caused by a higher amphiregulin secretion (31) into short, sharp pulses. This quick signal processing translates to a more ordered collective signalling without apoptosis (Fig. 3C). We also speculate that in this mutant, a mechanochemical feedback may explain the size of ERK waves. The PIK3CA^H1047R^ mutation increases cell motility (44, 45) as evidenced by angled traces in Fig. 3C, displacement of the centroid of collective events in Fig. S3, and cell movement in Video S4. The increased cell motility results in fluid-like epithelial deformations where cells frequently change their neighbours. It also alters the mechanochemical feedback between actomyosin cytoskeleton and ERK activity responsible for the propagation of ERK waves during epithelial wound repair (10). Consequently, large, long-lasting ERK waves result from largescale deformations of the epithelium.

Our quantification of the giant honeybee shimmering favours the “special-agents” hypothesis that suggest that groups of specialised bees initiate the shimmering (38, 42). Such groups also contribute to the spread of the wave-like process across the nest surface at speeds ranging up to 1 m/s which is much higher than expected by a mere bucket-bridging process (39). The experiments with dummy wasps recapitulate the real-world situation when wasps scan in front of the nest with predatory intent. However, experiments with free-flying real wasps allow measuring the critical distance in which the colony can respond by the anti-predator behaviour of shimmering (36). They also unravel another fascinating feature of this collective defence: the visual patterns of shimmering waves directionally align with the projected flight manoeuvres of the wasps preying in front of the bees’ nest (37). This phenomenon of predator driving was first reported in insects (37) and confirmed that shimmering honeybees utilise directional alignment to enforce their repelling power against preying wasps. Both capabilities of this shimmering behaviour, the extremely high speed of wave propagation, and the flexibility to quickly follow the predator in two dimensions with a visual signal, is only possible due to the sophisticated, logistical organisation between the leader and the follower. This coordination enables multiple initiations of the sequential wave event by saltatoric principles across the nest surface in any direction (24, 37, 38, 42).

### Using ARCOS

Our algorithm is versatile and therefore relies on several parameters tuned on a case-by-case basis. The analysis of time-lapse microscopy experiments requires image segmentation that yields a space-time point pattern with object coordinates and measurement changes over time, e.g., fluorescence intensity of a biomarker in single cells. The measurement has to be binarised to retain only active objects, which can be further processed by the spatial clustering and tracking algorithm in ARCOS. We suggested two binarisation approaches: one based on detrending, rescaling and thresholding of the measurement time series; a second based on local indicators of spatial association (LISAs) that quantify local measurement correlations that are then binarised. Binarisation involves parameters such as the size of the smoothing window for detrending, the neighbourhood radius for LISAs, and the threshold for final binarisation either of the rescaled measurement or the spatial correlation coefficient. The core ARCOS algorithm clusters active points in individual frames of the time-lapse and then tracks the clusters based on their overlap. The key parameters here are the neighbourhood radius and the minimal cluster size for spatial clustering. Supplementary R notebooks contain full analysis of all experimental modalities presented here and provide example parameters. The interactivity offered by our napari plugin is particularly useful when exploring datasets from 3D experiments such as the MCF10A acinus discussed above. For example, the interactive spaghetti plot lets browse the results of the analysis and highlight the events in the time-lapse movie.

### Concluding remarks

Our analysis demonstrated that single-cell signalling dynamics does not merely result from network wiring within individual cells, but also reflects emergent phenomena at the level of cell community. Disentangling these two contributions is necessary to further interpret oncogenic signalling. Future studies of spatio-temporal dynamics in other oncogenic MAPK/Akt pathway mutants will be essential to better characterise the dependency of individual cells on intercellular communication. Quantification of these complex emergent processes might then allow for modelling signalling networks in cell communities using agent-based approaches. This will enable true understanding of signalling processes at previously inaccessible length and time scales.

## Materials and Methods

### Cell culture

Wild-type human mammary MCF10A cells were a gift of J.S. Brugge, Harvard Medical School, Boston, MA. The isogenic variants of MCF10A harbouring the heterozygous mutations PIK3CA^H1047R^ and KRAS^G12V^ were a gift of Ben Ho Park, Vanderbilt University Medical Center, TN. NRK-52E expressing EKAREV-NLS were a gift from K. Aoki, National Institute for Basic Biology, Japan. Cultivation of wild-type MCF10A cells and the isogenic derivative was carried out in growth medium composed by DMEM:F12, 5% horse serum, 20 ng/ml recombinant human EGF (Peprotech), 10 *µ*g/ml insulin (Sigma-Aldrich/Merck), 0.5 *µ*g/ml hydrocortisone (Sigma-Aldrich/Merck), 200 U/ml penicillin and 200 *µ*g/ml streptomycin. NRK-52E cells were cultured in DMEM supplemented with 10% FBS, 200 U/ml penicillin and 200 mg/ml streptomycin.

### Biosensors and optogenetic actuator

To observe nuclei and measure ERK activity, we used the PiggyBac plasmid coding for the stable nuclear marker H2BmiRFP703 (pPBbSr2-H2B-miRFP703) and the PiggyBac plasmids coding for the ERK activity biosensor ERK-KTR fused with mRuby2 (pHygro-PB-ERK-KTR-mRuby2) or mTurquoise2 (pHygro-PB-ERK-KTR-mTurquoise2). To express the CRY2-based optogenetic actuator optoRAF and the plasma membrane anchor CIBN-KrasCT, we used a PiggyBac plasmid expressing both proteins separated by the selfcleaving peptide P2A (pPB3.0-PuroCRY2-cRAF-mCitrineP2A-CIBN-KrasCT). All plasmids were generated for a previous study from our laboratory (5).

To generate cells stably expressing the biosensors or the optogenetic actuator, we transfected the PiggyBac plasmids together with the helper plasmid expressing the transposase. The transfection of wild-type MCF10A cells and the mutated derivatives was carried out with FuGene (Promega) according to the manufacturer’s protocol. Antibiotic selection and image-based screening were used to generate stable clones with the desired biosensors/optogenetic tools.

### Live microscopy and optogenetics

MCF10A and NRK52E cells were seeded on 5 mg/ml fibronectin (PanReac AppliChem) on 24-well #1.5 glass bottom plates (Cellvis) to reach confluence at the day of the experiment. Image acquisition was executed with an epifluorescence Eclipse Ti inverted fluorescence microscope (Nikon) controlled by NISElements (Nikon) with a Plan Apo air 20x (NA 0.8) or Plan Apo air 40x (NA 0.9) objectives, and an Andor Zyla plus camera at a 16-bit depth. Laser-based autofocus was used throughout the experiments. Excitation and emission of the fluorescent protein was obtained with the following monochromatic LEDs and filters. Far red: 640 nm, ET705/72m; red: 555 nm, ET652/60m; NeonGreen: 508 nm, ET605/52; mTurquoise2: 440 nm, HQ480/40. For optogenetic experiments, cells expressing optoRAF were kept in the dark for at least 24h before the experiments and all preparatory steps at the microscope were carried out with red or green light (wavelength > 550 nm). Stimulation of the optoRAF was obtained with monochromatic 488 nm blue light for 100 ms at 0.3 W/cm2. The stimulated areas were produced by a digital mirror device (DMD; Andor Mosaic3) controlled by NIS Jobs based on masks prepared beforehand.

### Image analysis of 2D epithelia

We used two image segmentation approaches. In the first, we used Ilastik (46) to classify pixels in the nuclear channel and to build a nuclear probability mask. The classifier was trained on manual annotations of nuclei based on the H2B-mRFP703 fluorescent marker for MCF10A cells and EKAREV-NLS for NRK-52E. We used CellProfiler 3.0 (47) to perform further analysis. A threshold-based segmentation identified nuclei from the probability masks. The nuclear segmentation masks were used to quantify the average nuclear pixel intensity in the ERK-KTR channel. To determine the area corresponding to the cytosol, we expanded the nuclei by a predefined number of pixels, but not further than the underlying fluorescence intensity in the ERK-KTR channel. We calculated the average cytosolic pixel intensity of ERK-KTR from the resulting cytosolic ring. By dividing the average pixel intensities from the cytosolic ring and the nucleus, we obtained the cytosolic/nuclear ratio that estimates the ERK kinase activity. For FRET experiments with NRK-52E cells that express EKAREV-NLS biosensor, we calculated the FRET ratio, where the FRET image is divided pixel-by-pixel by the Donor image. Tracking of single cells was done on nuclear centroids in MATLAB using *µ*-track 2.2.1 (48).

In the second approach, for nuclear segmentation we used a custom segmentation algorithm that extracts image features from the nuclear marker channel using filters from a modified VGG16 (49) convolutional neural network (CNN) pretrained on ImageNet (50). We then quantified nuclear and cytosolic fluorescence intensity following the procedure described above. To track nuclei, we used a Python implementation of the linking algorithm (51), provided in the trackpy library (52).

Apoptotic events were manually annotated based on nuclear shrinkage, as reported earlier (5).

### Signal binarisation

Single-cell ERK activity obtained from image segmentation yields a (*x, y, m, t*) spatio-temporal point pattern, where (*x, y*) are centroid coordinates of nuclei, *m* is the corresponding measurement value at these coordinates (e.g., ERK activity), and *t* denotes time. To identify collective waves, we identify “active cells”, i.e., cells with *m* above a certain threshold. We implemented 3 methods in the `binMeas` function of the ARCOS R package to identify activity periods in time series. In the simplest case (the `biasMet` parameter set to `“none”`) the measurement is rescaled to [0, 1], smoothed with a short-term median filter to remove technical noise, and binarised with a global threshold. Setting `biasMet=“runmed”` activates detrending, where a long-term median-smoothed measurement is subtracted from the initial time series. With `biasMet=“lm”` the detrending step subtracts a linear fit to the measurement. An example of binarisation with the `runmed` method is shown in Fig. S1C. For the LISA-based activity binarisation, we used the statistics available in the R package elsa (34). These include local Moran’s *I* and local Geary’s *c* (27), as well as *G* and *G*^*∗*^ statistics (33). The resulting spatial correlation is binarised with a fixed threshold, as exemplified in Fig. S1D.

### Quantification of space-time correlation

In Figs. 2E, S2, and S4 we calculated a global Moran’s *I* autocorrelation coefficient according to the method described in (28). We applied the `Moran.I` function from the R package ape (53) to the (*x, y, m*) point pattern and calculated the coefficient independently for every time frame *t*. From the observed Moran’s *I* we subtracted the expected value of *I* under the null hypothesis that there was no correlation.

### MCF10A acini

Live imaging of 3D mammary acini with MCF10A cells and the corresponding image analysis were executed as described earlier (7). MCF10A cells were dissociated to single-cells and embedded in growth factor-reduced Matrigel (Corning) and overlaid with DMEM/F12 supplemented with 2% horse serum, 20 ng/ml recombinant human EGF, 0.5 mg/ml hydrocortisone, 10 mg/ml insulin, 200 U/ml penicillin and 200 mg/ml streptomycin. After 3 days, EGF, insulin and horse serum were removed from the medium. 25 mM Hepes was added to the medium before imaging.

Image acquisition was performed with an epifluorescence Eclipse Ti2 inverted fluorescence microscope (Nikon) equipped with a CSU-W1 spinning disk confocal system (Yokogawa) and a Plan Apo VC 60X water immersion objective (NA = 1.2). Images were acquired with a Prime 95B camera at 16 bit depth (Teledyne Photometrics). 3D segmentation of nuclei and extraction of cytosolic/nuclear ERKKTR fluorescence intensity ratio were performed using a customised version of the LEVER software (54).

### Shimmering bees

Image acquisition was performed as described earlier (35). To segment the act of flipping the abdomen in individual bees, we used a pixel classifier from the Ilastik image processing software (46). We obtained a probability mask that could be thresholded and processed as a conventional single-cell time-lapse movie that resulted in a single-bee time series data. Motion-active bees, i.e., bees that flipped their abdomen, were identified with the *G*^*∗*^ statistic, which we then thresholded and fed into ARCOS.

### Data & code availability

The data, R notebooks to reproduce the plots, and Jupyter notebooks to show the results in the napari image viewer were uploaded to the Mendeley Data repository (55). The code can be downloaded as: an R package (56), a python package (57), a napari plugin (58). A panel of synthetic tests can be executed with `devtools::test()` in the R console or with `pytest` in the command line. An extensive documentation of all packages can be accessed online (59). A video demonstration of the napari plugin from Video S9 is also available on YouTube (60).

## ACKNOWLEDGEMENTS

We would like to thank Prof. Yasuyuki Fujita and Yasuto Takeuchi for providing MDCK calcium imaging data. This work was supported by grants from the Swiss National Science Foundation (grants 31003A-163061 and 51PHPO-163583) and the Swiss Cancer League grant KLS-4867-08-2019 to Olivier Pertz, and from a Human Frontiers Science Program grant to Olivier Pertz and Andrew Cohen.

## AUTHOR CONTRIBUTIONS

Experiments: 2D epithelia – PAG, 3D acini – PE, giant honey bees – GK. BG developed the python package and the napari plugin. MD & M-AJ developed the R package. PAG, LH, MD, & AC performed image analysis. MD analysed and visualised the data. MD, PAG, & OP conceived the study and wrote the manuscript.

## Supplementary Figures

**Fig. S1.**
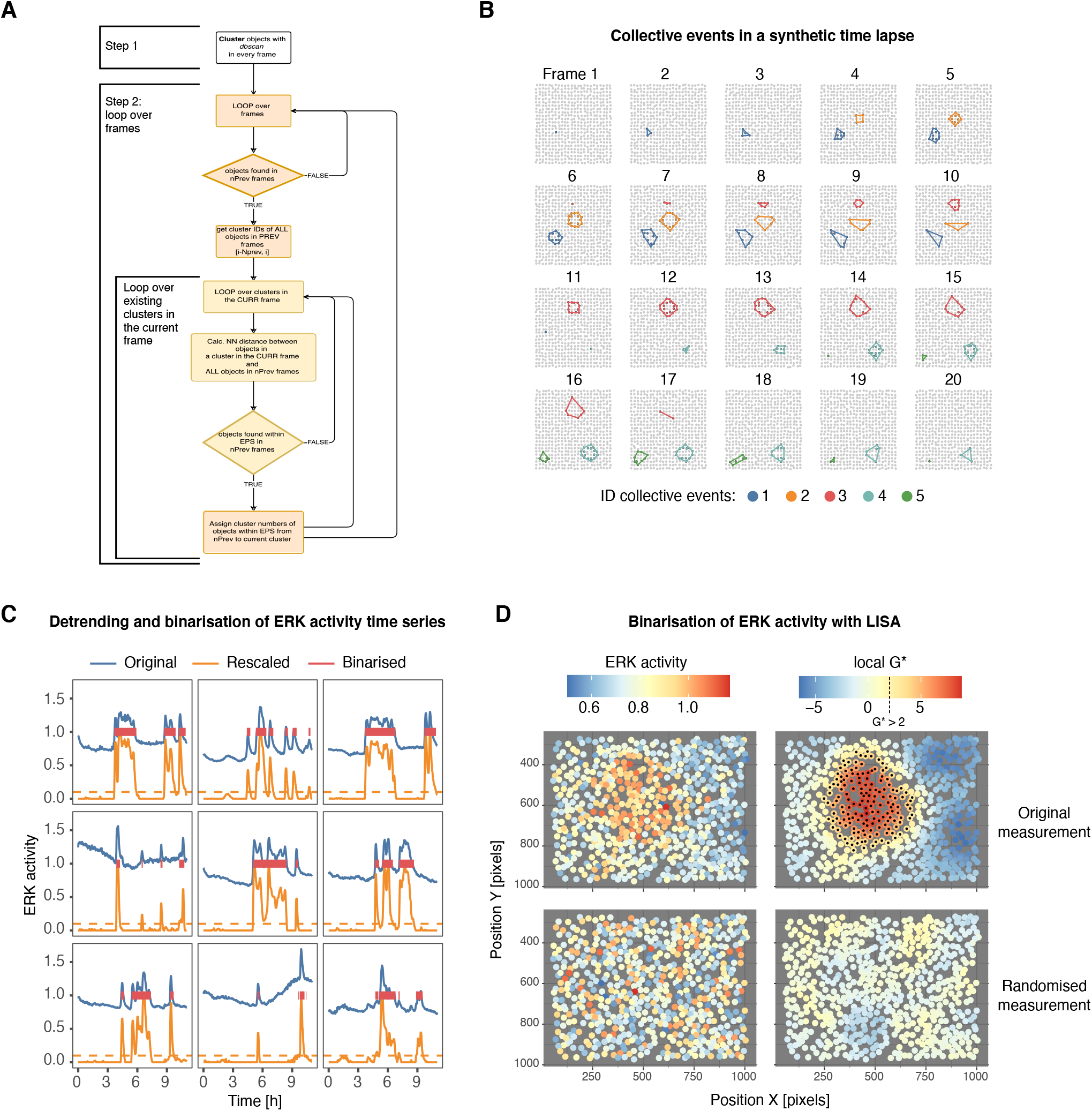
Algorithm outline, identification of collective events in synthetic data, and examples of measurement binarisation. (A) Flowchart of the core algorithm for detecting collective events in time-lapse data. (B) Synthetic, random time-lapse of collective activity events among cells positioned with a random jitter on a 32×32 grid. Each cell assumes only two states: active and inactive. Collective events detected by ARCOS are outlined by convex hulls. (C) Detrending and binarisation of single-cell ERK activity. Time series are first smoothed with a short-range, 5-time-point median filter. Then, the signal smoothed with a long-range, 501-time-point median filter is subtracted from the original signal. The result is rescaled between 0 and 1, and thresholded at 0.1 to binarise it. Red segments indicate periods of activity, which become an input for ARCOS. (D) Binarisation of ERK activity with a local indicator of spatial association (LISA). A sample frame with a pocket of cells with high ERK activity in the middle (upper row) and the same frame with the measurement randomised between cells while keeping the original X-Y positions (lower row). Cells with the local *G*^*∗*^ statistic greater than 2 are considered active (right column).

**Fig. S2.**
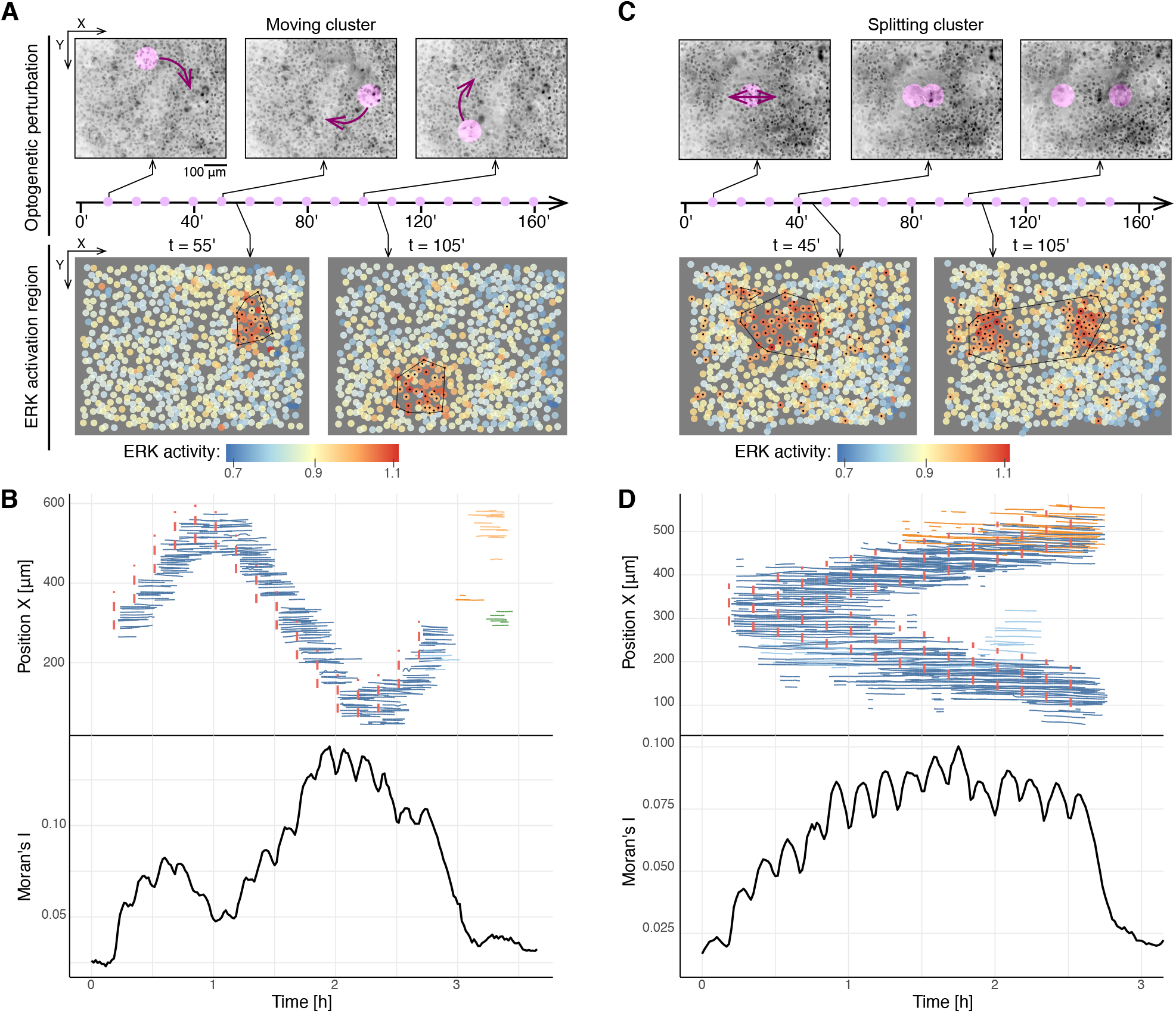
Identification of collective ERK signalling in MCF10A WT cells with optogenetically induced regions of ERK activity. (A) Repeated, pulsed optogenetic perturbation of a circular region moving clockwise along a circle. Upper row: inverted raw images from selected time points. Purple circles indicate the stimulation region. Purple dots on the timeline indicate the timing of a 100ms-long optogenetic pulse. Lower row: quantification of ERK activity from image segmentation for selected time points. Black dots indicate active cells calculated from binarisation; black polygons indicate the convex hull around collective events. (B) An X-axis projection of single cells from panel A during their participation in a collective event. Solid line colours indicate different collective events. Vertical dashed lines indicate the timing and position of the optogenetic stimulation. Black solid line is the Moran’s *I* measure of the spatial autocorrelation calculated from single-cell ERK activity data. (C,D) Same as A & B but the optogenetic perturbation is a circular region that splits horizontally over time.

**Fig. S3.**
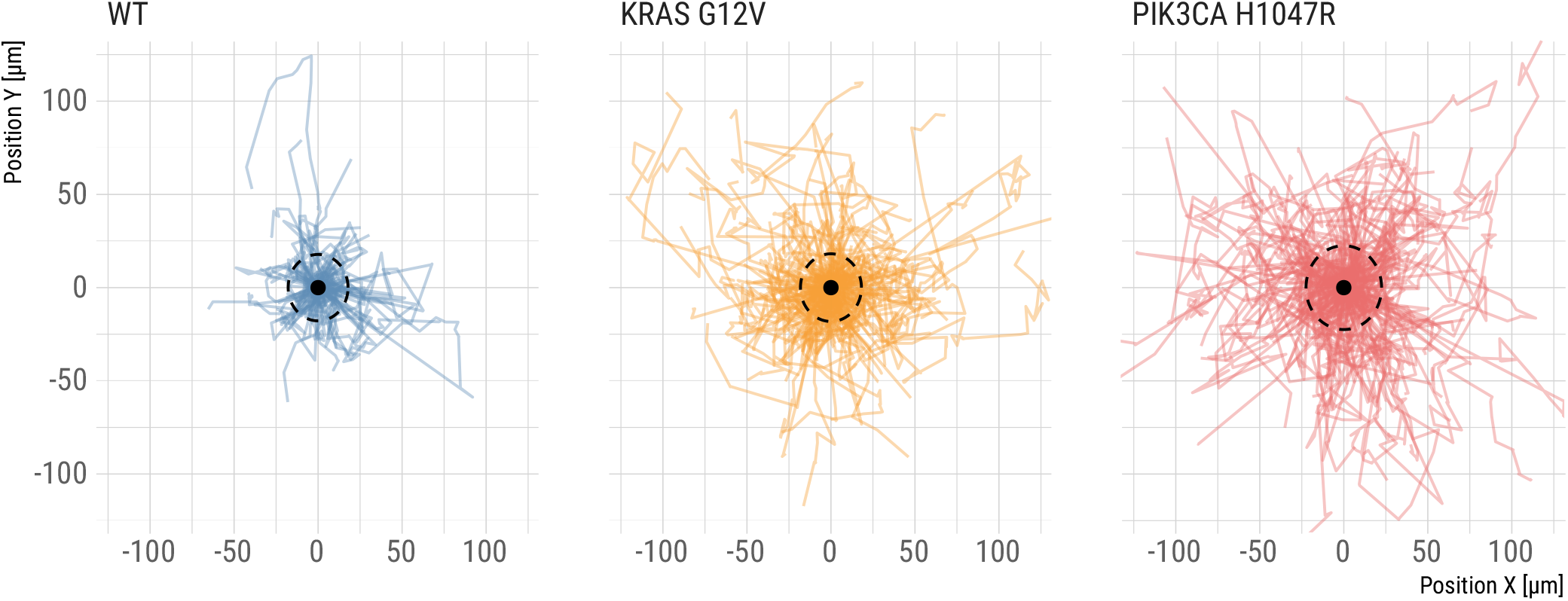
Displacement of collective events’ centroids in starved MCF10A WT, and KRAS^G12V^ and PIK3CA^H1047R^ mutant cell lines. The displacement for each collective event was calculated at every time frame based on the centroid of nuclei participating in the event. The origins of all displacements were moved to point (0,0) indicated with a black dot. Dashed circles indicate the time-averaged 1^st^ nearest neighbour radius for the respective cell line. A random sample of 200 events per cell line.

**Fig. S4.**
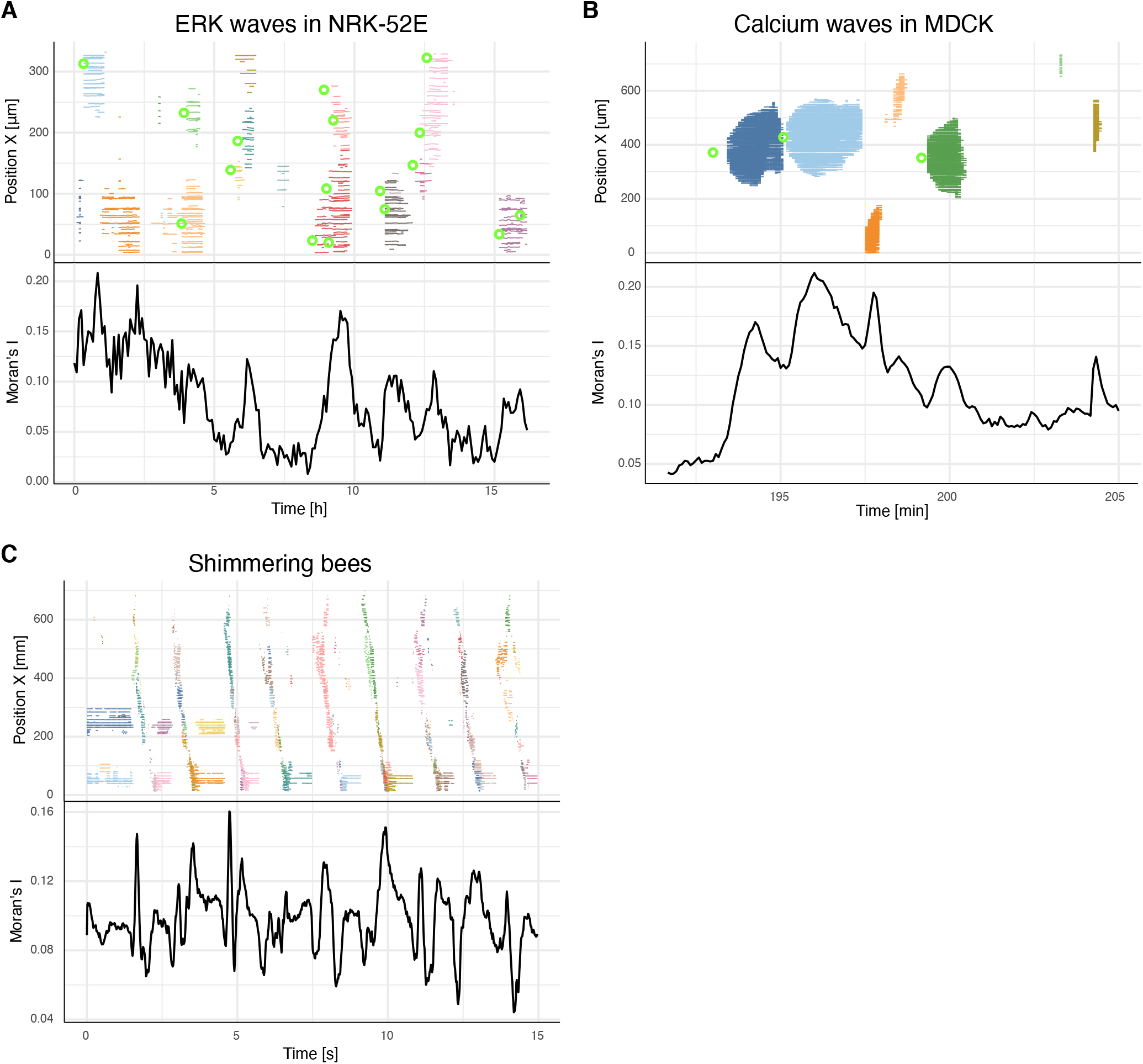
Spaghetti plots and quantification of the space-time correlation with a global indicator of spatial association, Moran’s *I* for ERK waves in NRK-52E epithelium (A), calcium waves in the MDCK epithelium (B), bee shimmering waves (C). Green rings indicate apoptotic events in panel A and apically extruded cells in panel B.

**Fig. S5.**
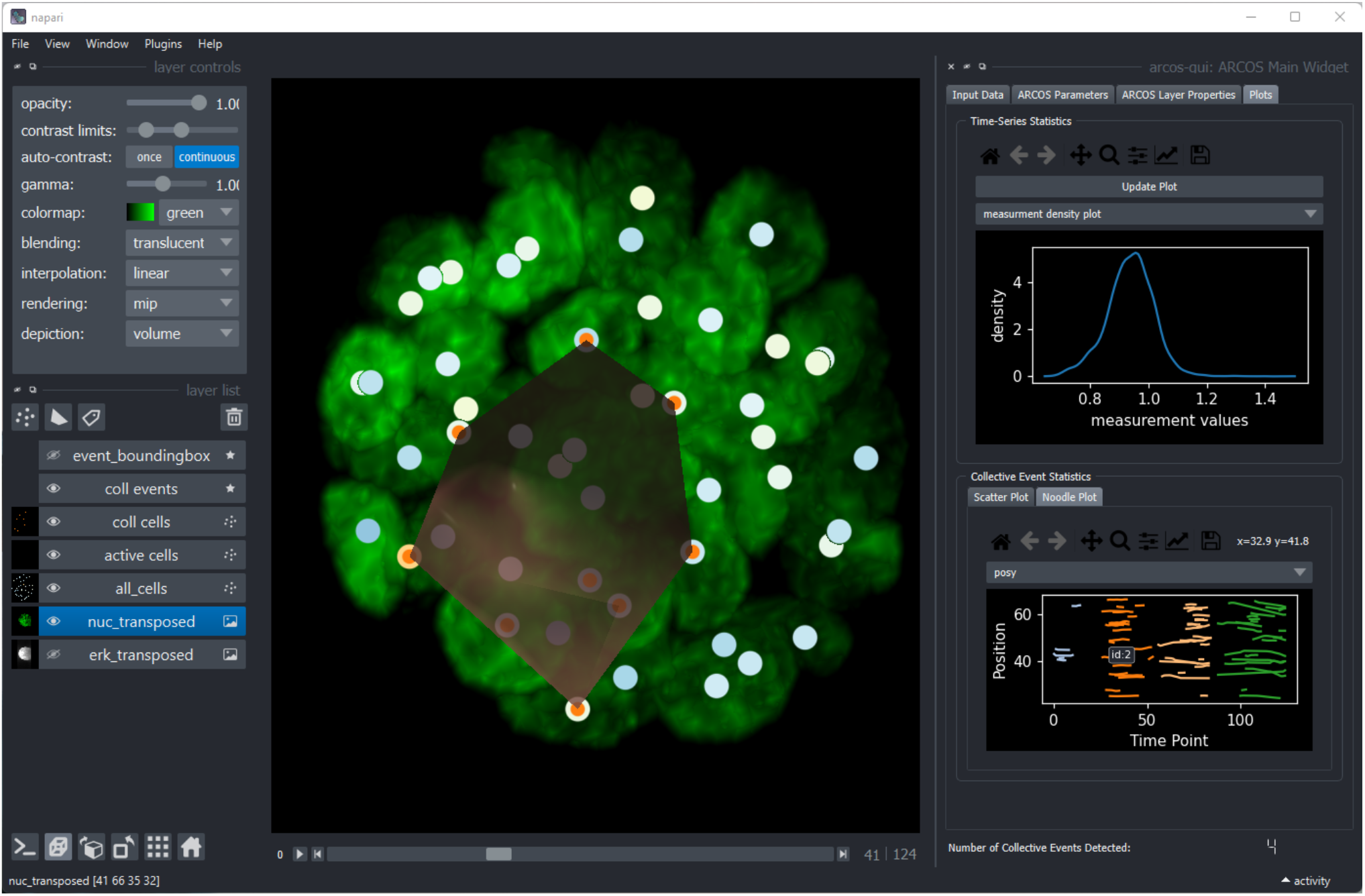
Quantification of collective ERK activation in a 7-day old MCF10A WT acinus. (A) Snapshots of a single collective event. Top row – maximum projection of the ERK-KTR channel; middle row – cell nuclei shaded according to detrended & normalised ERK activity; bottom row – nuclear centroids coloured by ERK activity and sized according to *z* distance. Black dots indicate active cells. Scale bar, 20 *µ*m. (B) A spaghetti plot of the space-time propagation of several collective events during the entire acquisition. The dashed rectangle indicates the space-time section from panel A.

## Supplementary Videos

**Video S1**. Starved MCF10A WT epithelial cells expressing the OptoRAF/ERK-KTR actuator/biosensor circuit are stimulated optogenetically with a growing circular region using DMD. Left panel – raw images from the ERK-KTR channel; right panel – nuclear centroids colour-coded according to the ERK-KTR cytoplasmic/nuclear ratio. Black dots – active cells identified in the binarisation step; shaded convex hulls – collective events identified by ARCOS. The stimulation region is indicated in magenta.

**Video S2**. Starved MCF10A WT epithelial cells expressing the OptoRAF/ERK-KTR actuator/biosensor circuit are stimulated optogenetically with a moving circular region using DMD. Left panel – raw images from the ERK-KTR channel; right panel – nuclear centroids colour-coded according to the ERK-KTR cytoplasmic/nuclear ratio. Black dots – active cells identified in the binarisation step; shaded convex hulls – collective events identified by ARCOS. The stimulation region is indicated in magenta.

**Video S3**. Starved MCF10A WT epithelial cells expressing the OptoRAF/ERK-KTR actuator/biosensor circuit are stimulated optogenetically with a splitting circular region using DMD. Left panel – raw images from the ERK-KTR channel; right panel – nuclear centroids colour-coded according to the ERK-KTR cytoplasmic/nuclear ratio. Black dots – active cells identified in the binarisation step; shaded convex hulls – collective events identified by ARCOS. The stimulation region is indicated in magenta.

**Video S4**. Collective ERK activity waves in starved MCF10A WT, KRAS^G12V^ and PIK3CA^H1047R^ epithelial cells expressing the ERK-KTR biosensor. Results of the ARCOS analysis are overlaid on raw images from the ERK-KTR channel. Each collective event is indicated with a distinct colour from a 9-colour palette, reused over time.

**Video S5**. Collective ERK activity waves in starved NRK-52E epithelial cells expressing the ERK activity reporter EKAREVNLS. The left panel depicts nuclei and the right panel nuclear centroids colour-coded according to the FRET ratio. Green circles – apoptotic cells; black dots – active cells identified in the binarisation step; shaded convex hulls – collective events identified by ARCOS.

**Video S6**. Collective calcium waves in starved MDCK epithelial cells expressing the calcium reporter GCaMP6S. Left panel – raw images from the GCaMP6S channel; right panel – nuclear centroids colour-coded according to the ERK-KTR cytoplasmic/nuclear ratio. Green circles – apically extruded cells; black dots – active cells identified in the binarisation step; shaded convex hulls – collective events identified by ARCOS.

**Video S7**. Bee shimmering in the giant honey bee colony. Left panel – raw images of the bee colony; right panel – bee centroids colour-coded according to the amount of shimmering. Black dots – “active bees” identified in the binarisation step; shaded convex hulls – collective events identified by ARCOS.

**Video S8**. Collective ERK activity waves in a stage 2 MCF10A WT acinus expressing the ERK-KTR biosensor. Left panel – maximum Z projection of the ERK-KTR channel; middle panel – 3D rendering of cell nuclei volumes with lever.js software, where gray shades indicate low/high ERK activities; right panel – nuclear centroids colour-coded according to collective events identified with ARCOS. The size of the nuclei in the right panel corresponds to their Z position.

**Video S9**. Video demonstration of features implemented in the interactive napari plugin. Demonstrates how to load and preprocess data, how to choose parameters to detect collective events and visualise them in interactive plots.

## Bibliography

1. Jeremy E. Purvis and Galit Lahav. Encoding and Decoding Cellular Information through Signaling Dynamics. Cell, 152(5):945–956, February 2013. ISSN 0092-8674. doi: 10.1016/j.cell.2013.02.005.

2. John G. Albeck, Gordon B. Mills, and Joan S. Brugge. Frequency-Modulated Pulses of ERK Activity Transmit Quantitative Proliferation Signals. Molecular Cell, 49(2):249–261, January 2013. ISSN 10972765. doi: 10.1016/j.molcel.2012.11.002.

3. Hyunryul Ryu, Minhwan Chung, Maciej Dobrzyński, Dirk Fey, Yannick Blum, et al. Frequency modulation of ERK activation dynamics rewires cell fate. Molecular Systems Biology, 11(11):838, November 2015. ISSN 1744-4292. doi: 10.15252/msb.20156458.

4. Timothy J. Aikin, Amy F. Peterson, Michael J. Pokrass, Helen R. Clark, and Sergi Regot. MAPK activity dynamics regulate non-cell autonomous effects of oncogene expression. eLife, 9:e60541, September 2020. ISSN 2050-084X. doi: 10.7554/eLife.60541.s

5. Paolo A. Gagliardi, Maciej Dobrzyński, Marc-Antoine Jacques, Coralie Dessauges, Pascal Ender, et al. Collective ERK/Akt activity waves orchestrate epithelial homeostasis by driving apoptosis-induced survival. Dev Cell, 56(12):1712–1726.e6, June 2021. ISSN 1878-1551. doi: 10.1016/j.devcel.2021.05.007.

6. Kelvin W. Pond, Olga Alkhimenok, Jayati Chakrabarti, Yana Zavros, Curtis A. Thorne, and Andrew L. Paek. Live-Cell Imaging in Human Colonic Monolayers Reveals Erk Waves Limit the Stem Cell Compartment to Maintain Epithelial Homeostasis. bioRxiv, February 2022. doi: 10.1101/2022.02.23.481374.

7. Pascal Ender, Paolo Armando Gagliardi, Maciej Dobrzyński, Coralie Dessauges, Thomas Höhener, et al. Spatio-temporal Control of ERK Pulse Frequency Coordinates Fate Decisions during Mammary Acinar Morphogenesis. bioRxiv, page 2020.11.20.387167, June 2022. doi: 10.1101/2020.11.20.387167.

8. Alessandro De Simone, Maya N. Evanitsky, Luke Hayden, Ben D. Cox, Julia Wang, et al. Control of osteoblast regeneration by a train of Erk activity waves. Nature, 590(7844):129–133, February 2021. ISSN 1476-4687. doi: 10.1038/s41586-020-03085-8.

9. Toru Hiratsuka, Yoshihisa Fujita, Honda Naoki, Kazuhiro Aoki, Yuji Kamioka, and Michiyuki Matsuda. Intercellular propagation of extracellular signal-regulated kinase activation revealed by in vivo imaging of mouse skin. eLife, 4:e05178, February 2015. ISSN 2050-084X. doi: 10.7554/eLife.05178.

10. Kazuhiro Aoki, Yohei Kondo, Honda Naoki, Toru Hiratsuka, Reina E. Itoh, and Michiyuki Matsuda. Propagating Wave of ERK Activation Orients Collective Cell Migration. Dev Cell, 43(3):305–317.e5, November 2017. ISSN 1878-1551. doi: 10.1016/j.devcel.2017.10.016.

11. Naoya Hino, Leone Rossetti, Ariadna Marín-Llauradó, Kazuhiro Aoki, Xavier Trepat, et al. ERK-Mediated Mechanochemical Waves Direct Collective Cell Polarization. Developmental Cell, 53(6):646–660.e8, June 2020. ISSN 1534-5807. doi: 10.1016/j.devcel.2020.05.011.

12. Yasuto Takeuchi, Rika Narumi, Ryutaro Akiyama, Elisa Vitiello, Takanobu Shirai, et al. Calcium Wave Promotes Cell Extrusion. Current Biology, 30(4):670–681.e6, February 2020. ISSN 0960-9822. doi: 10.1016/j.cub.2019.11.089.

13. Stefano Di Talia and Massimo Vergassola. Waves in Embryonic Development. Annu. Rev. Biophys., 51(1):annurev–biophys–111521–102500, May 2022. ISSN 1936-122X, 1936-1238. doi: 10.1146/annurev-biophys-111521-102500.

14. Roland Wedlich-Söldner and Timo Betz. Self-organization: the fundament of cell biology. Philos Trans R Soc Lond B Biol Sci, 373(1747):20170103, May 2018. ISSN 1471-2970. doi: 10.1098/rstb.2017.0103.

15. Alan L. Hodgkin, Andrew F. Huxley, and Bernard Katz. Measurement of current-voltage relations in the membrane of the giant axon of Loligo. J Physiol, 116(4):424–448, April 1952. ISSN 0022-3751. doi: 10.1113/jphysiol.1952.sp004716.

16. Jeremy B. Chang and James E. Ferrell. Mitotic trigger waves and the spatial coordination of the Xenopus cell cycle. Nature, 500(7464):603–607, August 2013. ISSN 1476-4687. doi: 10.1038/nature12321.

17. Fernanda Alcantara and Marilyn Monk. Signal Propagation during Aggregation in the Slime Mould Dictyostelium discoideum. Journal of General Microbiology, 85(2):321–334, December 1974. ISSN 0022-1287. doi: 10.1099/00221287-85-2-321.

18. Edwin Gilland, Andrew L. Miller, Eric Karplus, Robert Baker, and Sarah E. Webb. Imaging of multicellular large-scale rhythmic calcium waves during zebrafish gastrulation. Proc Natl Acad Sci U S A, 96(1):157–161, January 1999. ISSN 0027-8424. doi: 10.1073/pnas.96.1.157.

19. Lawrence D. Robb-Gaspers and Andrew P. Thomas. Coordination of Ca2+ signaling by intercellular propagation of Ca2+ waves in the intact liver. J Biol Chem, 270(14):8102–8107, April 1995. ISSN 0021-9258. doi: 10.1074/jbc.270.14.8102.

20. Won-Gyu Choi, Masatsugu Toyota, Su-Hwa Kim, Richard Hilleary, and Simon Gilroy. Salt stress-induced Ca2+ waves are associated with rapid, long-distance root-to-shoot signaling in plants. Proc Natl Acad Sci U S A, 111(17):6497–6502, April 2014. ISSN 1091-6490. doi: 10.1073/pnas.1319955111.

21. Denis Noble. A modification of the Hodgkin–Huxley equations applicable to Purkinje fibre action and pace-maker potentials. J Physiol, 160:317–352, February 1962. ISSN 0022-3751. doi: 10.1113/jphysiol.1962.sp006849.

22. Xavier Serra-Picamal, Vito Conte, Romaric Vincent, Ester Anon, Dhananjay T. Tambe, et al. Mechanical waves during tissue expansion. Nature Phys, 8(8):628–634, August 2012. ISSN 1745-2473, 1745-2481. doi: 10.1038/nphys2355.

23. Charisios D. Tsiairis and Alexander Aulehla. Self-Organization of Embryonic Genetic Oscillators into Spatiotemporal Wave Patterns. Cell, 164(4):656–667, February 2016. ISSN 1097-4172. doi: 10.1016/j.cell.2016.01.028.

24. Gerald Kastberger, Thomas Hoetzl, Michael Maurer, Ilse Kranner, Sara Weiss, and Frank Weihmann. Speeding up social waves. Propagation mechanisms of shimmering in giant honeybees. PLoS One, 9(1):e86315, 2014. ISSN 1932-6203. doi: 10.1371/journal.pone.0086315.

25. Nicholas Sofroniew, Talley Lambert, Kira Evans, Juan Nunez-Iglesias, Grzegorz Bokota, et al. napari/napari: 0.4.15rc1, March 2022.

26. Martin Ester, Hans-Peter Kriegel, Jörg Sander, and Xiaowei Xu. A density-based algorithm for discovering clusters in large spatial databases with noise. In KDD-96 Proceedings, pages 226–231. AAAI Press, 1996.

27. Luc Anselin. Local Indicators of Spatial Association-LISA. Geographical Analysis, 27(2): 93–115, April 1995. ISSN 00167363. doi: 10.1111/j.1538-4632.1995.tb00338.x.

28. John L. Gittleman and Mark Kot. Adaptation: Statistics and a Null Model for Estimating Phylogenetic Effects. Systematic Biology, 39(3):227–241, September 1990. ISSN 1063-5157. doi: 10.2307/2992183.

29. Minjmaa Minjgee, Mahmoud Toulany, Rainer Kehlbach, Klaudia Giehl, and H. Peter Rode-mann. K-RAS(V12) Induces Autocrine Production of EGFR Ligands and Mediates Radioresistance Through EGFR-Dependent Akt Signaling and Activation of DNA-PKcs. International Journal of Radiation Oncology*Biology*Physics, 81(5):1506–1514, December 2011. ISSN 03603016. doi: 10.1016/j.ijrobp.2011.05.057.

30. John P. Gustin, Bedri Karakas, Michele B. Weiss, Abde M. Abukhdeir, Josh Lauring, et al. Knockin of mutant PIK3CA activates multiple oncogenic pathways. Proc. Natl. Acad. Sci. U.S.A., 106(8):2835–2840, February 2009. ISSN 0027-8424, 1091-6490. doi: 10.1073/pnas.0813351106.

31. Christian D. Young, Lisa J. Zimmerman, Daisuke Hoshino, Luigi Formisano, Ariella B. Hanker, and et al. Activating PIK3CA Mutations Induce an Epidermal Growth Factor Receptor (EGFR)/Extracellular Signal-regulated Kinase (ERK) Paracrine Signaling Axis in Basallike Breast Cancer*. Molecular & Cellular Proteomics, 14(7):1959–1976, July 2015. ISSN 15359476. doi: 10.1074/mcp.M115.049783.

32. Kazuhiro Aoki, Yuka Kumagai, Atsuro Sakurai, Naoki Komatsu, Yoshihisa Fujita, et al. Stochastic ERK Activation Induced by Noise and Cell-to-Cell Propagation Regulates Cell Density-Dependent Proliferation. Molecular Cell, 52(4):529–540, November 2013. ISSN 1097-2765. doi: 10.1016/j.molcel.2013.09.015.

33. Arthur Getis and J. K. Ord. The Analysis of Spatial Association by Use of Distance Statistics. Geographical Analysis, 24(3):189–206, July 1992. ISSN 0016-7363. doi: 10.1111/j.1538-4632.1992.tb00261.x. Publisher: John Wiley & Sons, Ltd.

34. Babak Naimi, Nicholas A.S. Hamm, Thomas A. Groen, Andrew K. Skidmore, Albertus G. Toxopeus, and Sara Alibakhshi. ELSA: Entropy-based local indicator of spatial association. Spatial Statistics, 29:66–88, March 2019. ISSN 2211-6753. doi: 10.1016/j.spasta.2018.10.001.

35. Gerald Kastberger, Michael Maurer, Frank Weihmann, Matthias Ruether, Thomas Hoetzl, et al. Stereoscopic motion analysis in densely packed clusters: 3D analysis of the shimmering behaviour in Giant honey bees. Front Zool, 8(1):3, 2011. ISSN 1742-9994. doi: 10.1186/1742-9994-8-3.

36. Gerald Kastberger, Evelyn Schmelzer, and Ilse Kranner. Social Waves in Giant Honeybees Repel Hornets. PLoS ONE, 3(9):e3141, September 2008. ISSN 1932-6203. doi: 10.1371/journal.pone.0003141.

37. Gerald Kastberger, Frank Weihmann, Martina Zierler, and Thomas Hötzl. iant honeybees (Apis dorsata) mob wasps away from the nest by directed visual patterns. Naturwissenschaften, 101(11):861–873, November 2014. ISSN 0028-1042, 1432-1904. doi: 10.1007/s00114-014-1220-0.

38. Gerald Kastberger, Frank Weihmann, and Thomas Hoetzl. Complex social waves of giant honeybees provoked by a dummy wasp support the special-agent hypothesis. Communicative & Integrative Biology, 3(2):179–180, March 2010. ISSN 1942-0889. doi: 10.4161/cib.3.2.10809.

39. Gerald Kastberger, Frank Weihmann, Thomas Hoetzl, Sara E. Weiss, Michael Maurer, and Ilse Kranner. How to Join a Wave: Decision-Making Processes in Shimmering Behavior of Giant Honeybees (Apis dorsata). PLoS ONE, 7(5):e36736, May 2012. ISSN 1932-6203. doi: 10.1371/journal.pone.0036736.

40. Gerald Kastberger, Frank Weihmann, and Thomas Hoetzl. Social waves in giant honeybees (Apis dorsata) elicit nest vibrations. Naturwissenschaften, 100(7):595–609, July 2013. ISSN 0028-1042, 1432-1904. doi: 10.1007/s00114-013-1056-z.

41. Sarah E. Radloff, H. R. Hepburn, and Michael S. Engel. The Asian Species of Apis. In H. Randall Hepburn and Sarah E. Radloff, editors, Honeybees of Asia, pages 1–22. Springer Berlin Heidelberg, Berlin, Heidelberg, 2011. ISBN 978-3-642-16421-7 978-3-642-16422-4. doi: 10.1007/978-3-642-16422-4_1.

42. Evelyn Schmelzer and Gerald Kastberger. ‘Special agents’ trigger social waves in giant honeybees (Apis dorsata). Naturwissenschaften, 96(12):1431–1441, December 2009. ISSN 0028-1042, 1432-1904. doi: 10.1007/s00114-009-0605-y.

43. Taryn E. Gillies, Michael Pargett, Marta Minguet, Alex E. Davies, and John G. Albeck. Linear Integration of ERK Activity Predominates over Persistence Detection in Fra-1 Regulation. Cell Systems, 5(6):549–563.e5, December 2017. ISSN 24054712. doi: 10.1016/j.cels.2017.10.019.

44. Huan Pang, Rory Flinn, Antonia Patsialou, Jeffrey Wyckoff, Evanthia T. Roussos, et al. Differential Enhancement of Breast Cancer Cell Motility and Metastasis by Helical and Kinase Domain Mutations of Class IA Phosphoinositide 3-Kinase. Cancer Res, 69(23):8868–8876, December 2009. ISSN 0008-5472, 1538-7445. doi: 10.1158/0008-5472.CAN-09-1968.

45. Yardena Samuels, Luis A. Diaz, Oleg Schmidt-Kittler, Jordan M. Cummins, Laura Delong, et al. Mutant PIK3CA promotes cell growth and invasion of human cancer cells. Cancer Cell, 7(6):561–573, June 2005. ISSN 1535-6108. doi: 10.1016/j.ccr.2005.05.014.

46. Stuart Berg, Dominik Kutra, Thorben Kroeger, Christoph N. Straehle, Bernhard X. Kausler, et al. ilastik: interactive machine learning for (bio)image analysis. Nat Methods, 16(12): 1226–1232, December 2019. ISSN 1548-7105. doi: 10.1038/s41592-019-0582-9.

47. Claire McQuin, Allen Goodman, Vasiliy Chernyshev, Lee Kamentsky, Beth A. Cimini, et al. CellProfiler 3.0: Next-generation image processing for biology. PLoS Biol, 16(7):e2005970, July 2018. ISSN 1545-7885. doi: 10.1371/journal.pbio.2005970.

48. Khuloud Jaqaman, Dinah Loerke, Marcel Mettlen, Hirotaka Kuwata, Sergio Grinstein, et al. Robust single-particle tracking in live-cell time-lapse sequences. Nat Methods, 5(8):695–702, August 2008. ISSN 1548-7105. doi: 10.1038/nmeth.1237.

49. Dewa Made Sri Arsa and Anak Agung Ngurah Hary Susila. VGG16 in Batik Classification based on Random Forest. In 2019 International Conference on Information Management and Technology (ICIMTech), pages 295–299, Jakarta/Bali, Indonesia, August 2019. IEEE. ISBN 978-1-72813-333-1. doi: 10.1109/ICIMTech.2019.8843844.

50. Jia Deng, Wei Dong, Richard Socher, Li-Jia Li, Kai Li, and Li Fei-Fei. ImageNet: A largescale hierarchical image database. In 2009 IEEE Conference on Computer Vision and Pattern Recognition, pages 248–255, Miami, FL, June 2009. IEEE. ISBN 978-1-4244-3992-8. doi: 10.1109/CVPR.2009.5206848.

51. John C. Crocker and David G. Grier. Methods of Digital Video Microscopy for Colloidal Studies. Journal of Colloid and Interface Science, 179(1):298–310, April 1996. ISSN 00219797. doi: 10.1006/jcis.1996.0217.

52. Daniel B. Allan, Thomas Caswell, Nathan C. Keim, Casper M. van der Wel, and Ruben W. Verweij. soft-matter/trackpy: Trackpy v0.5.0, April 2021.

53. Emmanuel Paradis and Klaus Schliep. ape 5.0: an environment for modern phylogenetics and evolutionary analyses in R. Bioinformatics, 35(3):526–528, February 2019. ISSN 1367-4803. doi: 10.1093/bioinformatics/bty633.

54. Mark Winter, Walter Mankowski, Eric Wait, Sally Temple, and Andrew R. Cohen. LEVER: software tools for segmentation, tracking and lineaging of proliferating cells. Bioinformatics, 32(22):3530–3531, November 2016. ISSN 1367-4811. doi: 10.1093/bioinformatics/btw406.

55. Maciej Dobrzyński. Source Data and R/Python Scripts. https://data.mendeley.com/datasets/8gcncg6zkt, 2022.

56. Maciej Dobrzyński and Marc-Antoine Jacques. ARCOS R Source Code. https://github.com/pertzlab/ARCOS, 2022.

57. Benjamin Grädel. ARCOS Python Source Code. https://github.com/pertzlab/arcos4py, 2022.

58. Benjamin Grädel. ARCOS napari GUI Source Code. https://github.com/pertzlab/arcos-gui, 2022.

59. Benjamin Grädel. ARCOS documentations. https://arcos.gitbook.io, 2022.

60. Benjamin Grädel. ARCOS GUI Demo. https://youtu.be/hG_z_BFcAiQ, 2022.

